# Dual-action nanoconjugate for overcoming r-tPA-resistant clots

**DOI:** 10.64898/2026.05.13.725039

**Authors:** Audrey Picot, Mandy Leboucher, Charly Helaine, Ankita Talukdar, Igor Khalin, Sara Martinez de Lizarrondo, Maxime Gauberti, Mialitiana Solo Nomenjanahary, Didier Goux, Benoit Ho-Tin-Noé, Denis Vivien, Thomas Bonnard

**Affiliations:** Normandie Université, UNICAEN, INSERM, PhIND (Physiopathology and Imaging of Neurological Disorders) Institut Blood, Brain & Memory @ Caen-Normandie, Cyceron, Caen, France; CHU CAEN, Radiology departement, CHU de Caen côte de Nacre, Caen, France; Optimisation Thérapeutique en Neuropsychopharmacologie, Université Paris Cité, U1144 Institut National de la Santé et de la Recherche Médicale (INSERM), Paris, France; Normandie Université, UNICAEN, CMAbio3: Centre de Microscopie Appliquée à la Biologie, US Emerode, 14000 Caen, France; CHU CAEN, Clinical Research Department, CHU de Caen côte de Nacre, Caen, France; Institute for Stroke and Dementia Research (ISD), LMU University Hospital, LMU Munich, Germany

## Abstract

Clot resistance to pharmacological thrombolysis remains a critical challenge in ischemic stroke (IS) management. Thrombus heterogeneity, particularly the presence of thrombolysis-resistant domains composed of dense fibrin and non-fibrin components, including neutrophil extracellular traps (NETs), significantly limits the efficacy of recombinant tissue-type plasminogen activator (r-tPA) and its variant, Tenecteplase (TNK). Consequently, novel therapeutic strategies are urgently required. Emerging evidence suggests that co-administration of deoxyribonuclease I (DNase I) with r-tPA can degrade DNA fibers and enhance clot lysis. In this study, we optimized a previously developed theranostic agent—iron oxide microparticles coated with polydopamine—by dual-grafting both r-tPA and DNase to target resistant thrombi. Using functional ultrasound imaging (fUS) during the acute phase of IS, we demonstrated accelerated reperfusion with this dual-functionalized platform in a r-tPA resistant IS model. Furthermore, MRI analysis confirmed a significant reduction in lesion volume at 24 hours, correlating with improved functional recovery five days post-ischemia.

## Introduction

Ischemic stroke (IS) remains the second leading cause of death and dementia worldwide (WSO). Despite extensive research and the development of new treatment strategies, only a limited number of pharmacological treatments are currently approved, namely recombinant tissue-type plasminogen activator (r-tPA, Alteplase®) and, more recently, its genetically modified variant, Tenecteplase (TNK, Metalyse®)^1,2^. However, the clinical use of these thrombolytics is restricted by their narrow therapeutic window (4.5 h for r-tPA) and their numerous contraindications, resulting in fewer than 20% of patients being eligible for treatment^3,4^. Moreover, even among treated patients, a large proportion fails to achieve complete or sustained recanalization, which severely limits therapeutic success.

This limited efficacy highlights the need to better understand the mechanisms underlying treatment failure. Increasing evidence suggests that one major contributor is clot resistance to fibrinolysis^5^. Several studies have demonstrated significant heterogeneity in thrombus composition among stroke patients, including fibrin-rich clots surrounded by a dense outer shell composed of platelets, dense fibrin, van Willebrand factor, neutrophils, and neutrophil extracellular traps (NETs)^6–8^. This dense structure forms a physical and biochemical barrier that limits the penetration of plasminogen and r-tPA, thereby impairing fibrinolysis. In addition, the presence of cross-linked fibrin and platelet aggregates further reinforces clot rigidity and stability, making them highly resistant to enzymatic degradation by r-tPA alone^6–8^.

To overcome these limitations, adjunctive or combinatorial approaches have been explored to enhance r-tPA efficacy, including the use of agents targeting non-fibrin components such as DNA or NETs. In particular, deoxyribonuclease I (DNase I) has shown the ability to degrade the extracellular DNA network within thrombi, thus loosening their structure and promoting fibrinolysis^9–11^. Nevertheless, the systemic administration of these enzymes is limited by their short plasma half-lives, lack of targeted delivery, and potential off-target effects.

In this context, the development of a nanoconjugate co-delivering r-tPA and DNase I directly to the clot surface represents a promising strategy to overcome fibrinolytic resistance. This platform, based on polydopamine coated particles, allows efficient enzyme immobilization and localized activity at the clot interface thereby enhancing thrombolysis while minimizing systemic side effects.

We aimed to design a multifunctional nanoconjugate capable of efficiently degrading r-tPA–resistant thrombi by simultaneously targeting their fibrin and extracellular DNA/NETs components. To this end, we developed iron oxide (IO) microparticles coated with polydopamine (PDA), a biocompatible polymer that provides versatile chemical groups for enzyme conjugation and surface stability^12^. The resulting nanoplatform, termed IO@PDA@tPA@DNase, was engineered to deliver r-tPA and DNase I in close proximity at the clot site. PDA shell ensures strong and stable immobilization of both enzymes through catechol–amine interactions. This dual-conjugation strategy preserves their enzymatic activity and allows a synergistic fibrinolytic effect, where DNase I first dismantles the DNA/NET shell, thereby facilitating deeper r-tPA diffusion and enhanced fibrin degradation.

## Material and Methods

### Conjugate comprising the particle IO@PDA, r-tPA and DNase I (Pulmozyne®)

The IO@PDA synthesis and the r-tPA conjugation was made as described before ^13^, except for the incubation time of the r-tPA which was reduced to 30 min. After removing the supernatant and measure the r-tPA quantity fixed on the IO@PDA, 500µl of DNase I (1,17 mg/ml, PULMOZYME 2500 U/2,5 ml sol p inhal p nébulis) was added to the particles and incubated 30 minutes at 4°C under constant agitation. After 30 minutes, a magnet is used to collect only the IO@PDA@tPA conjugated with DNase I that will be resuspended in a 0.3 M mannitol solution. Before discarding the supernatant, western blotting was used to determine the quantity of r-tPA and DNase I fixed on the particles.

### Western Blotting

To determine the concentration of unbound r-tPA and DNase I in the sample, a Western blot was performed using a standard curve of increasing concentrations of free r-tPA and DNase I. Samples were prepared and loaded onto stain-free precast gels (10% Mini-PROTEAN® TGX™ Protein Gels, 4561031, Bio-Rad) along with the r-tPA or DNase I standards. Proteins were visualized using stain-free technology (ChemiDoc, Bio-Rad) and quantified via densitometric interpolation using Image Lab software.

### Scanning electron microscopy (SEM)

IO@PDA and IO@PDA@tPA@DNase were dried, coated with platinum and characterized by a JSM-7200F Scanning Electron Microscope (JEOL Europe SAS, Croissy-sur-Seine, France) at 3 kV.

### Agarose gel electrophoresis for DNase I activity assay

To assess the nuclease activity of free and particle-bound DNase I, DNA degradation assays were performed using salmon sperm DNA (Invitrogen, ThermoFisher Scientific, 15632-011) as a substrate. DNA samples (0,1 µg/µL) were incubated at 37 °C for 30 minutes under the following conditions: (1) DNA alone (negative control), (2) DNA with free DNase I (1 µg or 2 µg), (3) DNA with IO@PDA particles (particle control), and (4) DNA with IO@PDA@tPA@DNase particles (1 µg or 2 µg). The final volume of each reaction was adjusted to 15 µL with nuclease-free water.

Following incubation, samples were mixed with loading buffer and loaded onto a 1% agarose gel containing a nucleic acid stain (GelRed Nucleic Acid Gel Stain, Fisher Scientific, NC9938951). Electrophoresis was carried out at 100 V for 30–40 minutes in 1X TAE buffer. DNA bands were visualized using a UV transilluminator or gel imaging system (Bio-Rad). The degradation of salmon DNA was assessed qualitatively by comparing the band intensity and pattern of DNA bands across conditions, confirming whether the enzyme retained its activity when conjugated to particles.

### Clot degradation assay

For thrombus collection, patients were treated in Rothschild Foundation hospital by EVT (endovascular therapy) with successful thrombi retrieval. The EVT procedure was chosen at the interventionalist’s discretion using a stent-retriever and/or a contact aspiration technique. Thrombi were collected at the end of EVT (Endovascular Treatment in Ischemic Stroke registry, URL: https://www.clinicaltrials.gov; Unique identifier: NCT03776877) and were preserved at -80°C. For the clot degradation assay, each clot was bisected to allow comparison of two treatment conditions using matched clot material. Clot halves were transferred into individual wells of a 24-well plate and incubated in 150 µL of BAPA (benzylsulfonyl D argininyl prolyl 4 amidinobenzylamide) plasma (Hirudin-based anticoagulated plasma) for 1 hour at 37 °C with gentle shaking (400 RPM) to allow rehydration. After rehydration, clots were gently blotted to remove excess fluid and weighed individually using an analytical balance to determine baseline mass. Treatments were then applied directly onto the clots. Treatment solutions were prepared in HBSS (Hank’s Balanced Salt Solution) containing Ca²⁺ and Mg²⁺, supplemented with BAPA plasma. The total volume of treatment applied to each clot was adjusted based on clot weight, using a ratio of 40 µL per mg of clot. Final concentrations were 1 µg/mL for tissue plasminogen activator (r-tPA) and 0.6 µg/mL for DNase I when applicable. Plates were incubated at 37 °C with shaking (400 RPM) for 2 hours. Clots were removed and weighed every 20 minutes to monitor time-dependent degradation. Three pairwise comparisons were performed: (i) r-tPA vs IO@PDA@tPA; (ii) r-tPA + DNase I vs IO@PDA@tPA@DNase; (iii) IO@PDA@tPA vs IO@PDA@tPA@DNase. Each clot was used for a single treatment comparison, ensuring intra-clot consistency between conditions (n=15). Lastly, an overall comparison was conducted for descriptive purposes only.

### *In vivo* experiments Animals

All experiments were conducted in compliance with French ethical law (Decrees 2013-118 and 2020-274) and the European Communities Council guidelines (2010/63/EU). Experiments were approved by the local ethical committee of Normandy (CENOMEXA, APAFIS #45085). Animals were provided and maintained under specific pathogen-free conditions at the Centre Universitaire de Ressources Biologiques (CURB, Basse-Normandie, France), and all had free access to food and tap water. Mice were housed in a temperature-controlled room on a 12-hour light/12-hour dark cycle with food and water *ad libitum*. Animals were randomly assigned to the different experimental groups, and all experiments and subsequent analyses were performed in a blinded manner. A catheter was inserted into the tail vein of the mice for intravenous administration of the conjugate or treatments before the AlCl_3_ IS model and Ultrasound (fUS) imaging acquisition. After surgery, animals were allowed to recover in a clean heated cage before taking them back to the animal facility.

### AlCl_3_ ischemic stroke model

Mice were anesthetized with isoflurane (5%, 70/30 NO_2_/O_2_). Mice were placed in a stereotaxic device and maintained under anaesthesia with isoflurane (1.5-2%, 70/30 NO_2_/O_2_) at 37°C by the integrated heat animal holder. Before beginning the surgery, buprenorphine (BUPRECARE, H0270. 54000561, 0.3mg/ml) was injected to the mice for analgesia. The MCA was exposed and a piece of AlCl_3_ (Sigma-Aldrich) -saturated filter paper was topically applied on the artery for 5 minutes.

### Ultrasound (fUS) imaging

All ultrasound imaging acquisitions were performed from bregma -3 mm to bregma +1 mm, both before and 15 minutes after stroke onset, using an ultrafast scanner (Iconeus One, Iconeus, France), the IcoScan acquisition software, and a dedicated ultrasound probe (Iconeus IcoPrime-4D MultiArray, 15 MHz, 256 elements, 100 µm pitch). Mice were placed in a stereotaxic frame. After incision of the scalp over the skull, both temporal crests were thinned to facilitate ultrasound wave penetration into the cortex and to prevent resolution loss. A cranial window was performed only on the left side to access the MCA (to avoid signal variations due to drilling). The skull was then cleaned with saline. Centrifuged ultrasound gel (2,500 × g, 5 min) was applied to the skull before placing the probe. ***CBV monitoring:*** A 5-minute acquisition was performed before stroke induction to obtain baseline CBV (T0). A 45-minute acquisition was then conducted 15 minutes after stroke onset to monitor CBV variations. Five minutes after the start of the 45-minute acquisition (i.e., 20 minutes after stroke onset), mice received one of the following treatments: Saline; IO@PDA (2 mg.kg); IO@PDA@tPA@DNase (2.5 mg/kg r-tPA equivalency and 1.32 mg/kg DNase equivalency); IO@PDA (2 mg/kg) + r-tPA (2.5 mg/kg) + DNase (1.32 mg/kg); DNase (1.32 mg/kg) or r-tPA (10 mg/kg)). ***CBV variations:*** Two brain regions were selected for analysis: the right isocortex corresponding to the contralateral hemisphere and the ipsilateral hemisphere corresponding to the lesion site. Average CBV in each area was calculated over the 5-minute baseline acquisition and at each 5-minute interval during the post-ischemic acquisition. To evaluate reperfusion in the ipsilateral cortex following treatment, CBV was further analysed every 10 minutes. CBV variations were expressed as the percentage change relative to the baseline CBV.

### MRI acquisition

Experiments were performed using a BioSpec 7-T TEP-MRI system with a surface coil resonator (Bruker, Germany). Mice were anesthetized with isoflurane (1.5 to 2.0%) and maintained at 37°C by the integrated heat animal holder, and the breathing rate was monitored during the imaging procedure.

Brain scans including of *T_2_*-weighted (RARE sequence, with TR/TE = 3500 ms/40 ms), *T \**-weighted sequences (fast-low angle shot (FLASH) sequence, with TR/TE = 50 ms/3.5 ms) and TOF sequences (TR/TE = 12 ms/4.2 ms) were made 24 hours after treatments injection to visualize the lesion size, hemorrhagic transformation and the recanalization.

### Corridor test

To assess sensorimotor function, we choose a homemade task which is based on the ability of animals to explore objects. The protocol was previously described in^13^.

### Corner test

The corner test was used to assess sensorimotor and post-stroke asymmetry. Mice were placed between two angled vertical boards forming a 30° corner. When entering the corner, mice typically rear and turn either to the left or the right to exit. Each mouse underwent 20 consecutive trials per session. The direction of turning (left or right) was recorded for each trial. A preference for turning contralateral to the lesion side was interpreted as sensorimotor impairment. The test was conducted before, 24h and 5 days after stroke onset. Results are presented as the percentage of turns toward each side, indicating turning side preference.

### Immunostaining

Five minutes post-treatment administration and lectin injection (only for two-photon microscopy), corresponding to 25 minutes post IS, mice were euthanized for brain collection. Harvested tissues were immediately fixed in 4% paraformaldehyde (PFA) for 24 hours at 4°C followed by two successive 24-hour washes in PBS sucrose (20%) for cryoprotection.

For two-photon microscopy: Fixed brains were sectioned at 80 µm using a vibratome (Leica Biosystems VT 1000S, Wetzlar, Germany). Sections were permeabilized in PBS containing 0.1% Tween-20 for 60 minutes at room temperature (RT) under agitation. Non-specific binding was blocked for 120 minutes in a blocking buffer (PBS, 1% BSA, 10% donkey serum, and 0.5% Triton X-100). Sections were then incubated with primary antibodies against Fibrinogen (1:1000; Stago, #IGYFI100) and CD42b (1:800; Invitrogen, #14-0429-82) in blocking solution for 72 hours at 4°C. After three 10-minute washes in PBS, samples were incubated for 3 hours at RT with Cy5- and Cy3-conjugated AffiniPure F(ab’)2 fragments of donkey IgG (1:800; Jackson ImmunoResearch). Following three final washes, sections were mounted on Superfrost™ Plus slides using Fluoromount-G™. Imaging was performed using a Bruker Ultima 2P Plus microscope (Billerica, MA, USA) equipped with an Olympus XLUMPlanFL N 20×/1.0 W objective. Excitation was provided by a Chameleon Vision II laser (Coherent, Glasgow, Scotland) tuned to 920 nm (power: 300 Pockels). Signal was collected via PMT detectors (Master Gain: 500) through red, green, and blue filters. Z-stacks were acquired at a depth of 50 μm (Galvo mode, dwell time 3.6 µs, 1024 × 1024 or 2048 × 2048 pixels, 13-bit).

For Epifluorescence microscopy: Fixed brains were embedded in Cryomatrix™ and frozen at -80°C. Coronal sections (10 µm) were cut using a cryostat (Leica CM3050S) and mounted on Superfrost™ Plus adhesive slides. After rehydration in PBS (3 × 15 min), sections were incubated overnight at room temperature with primary antibodies against Fibrinogen (1:1000; Stago, #IGYFI100) and CD42b (1:800; Invitrogen, #14-0429-82). Following three PBS washes, AffiniPure F(ab’)2 fragments of donkey IgG conjugated to Cy3 (for Fibrinogen) and Cy5 (for CD42b) (1:800; Jackson ImmunoResearch) were applied for 2 hours at room temperature. Slides were then mounted using Fluoromount™ medium containing DAPI for nuclear counterstaining. Images were acquired using a Leica DM6000B upright epifluorescence microscope equipped with a Hamamatsu Orca Flash 4.0 camera and MetaMorph software (Version 7.8.13.0). Image analysis and processing were performed using ImageJ (NIH).

### Statistical analysis

All results are presented as mean ± Standard Deviation (SD). Statistical analyses were performed blinded to the experimental groups, using Graph Pad Prism software (v8.0). We assumed normality of the data distribution with Shapiro-Wilk test. Two-way ANOVA was used for comparing more than two groups. Differences were considered statistically significant if p-value < 0.05.

### Acceptance Criteria

Mice were excluded in case of death during the experimental procedure, technical problems, reperfusion before treatment administration or hemorrhage during the surgery.

## Results

Building on our prior development of a nanoconjugate consisting of iron oxide nanoparticles surrounded by polydopamine and functionalized with r-tPA^13^, we thought to improve IS treatment by specifically addressing r-tPA blood clot resistance. Thus, a new dual conjugate, IO@PDA@tPA@DNase, was designed. Its synthesis followed the previously described protocol for IO@PDA formation^12^, while the sequential conjugation of r-tPA^13^ and DNase I was carried out using similar approach (Figure 1A); grafting via primary amine to polydopamine surface via Michael’s addition. The amount of r-tPA grafted on the surface of the IO@PDA remains consistent with previous report^13^ (0.55 mg/mL; Table 1, Figure 1B). The quantity of DNase I, assessed by western blotting, conjugated to the particles is around 0.3 mg/mL (Table 1, Figure 1C). Scanning electron microscopy (SEM) images demonstrate that the dual grafting of r-tPA and DNase onto the IO@PDA surface does not alter their structural morphology (Figure 1D, E). Moreover, the particle sizes observed via SEM are consistent with the hydrodynamic diameters previously measured (614 nm for bare particles and 660 nm for the nanoconjugates; Table 1, Figure 1D, E).

**Figure 1.**
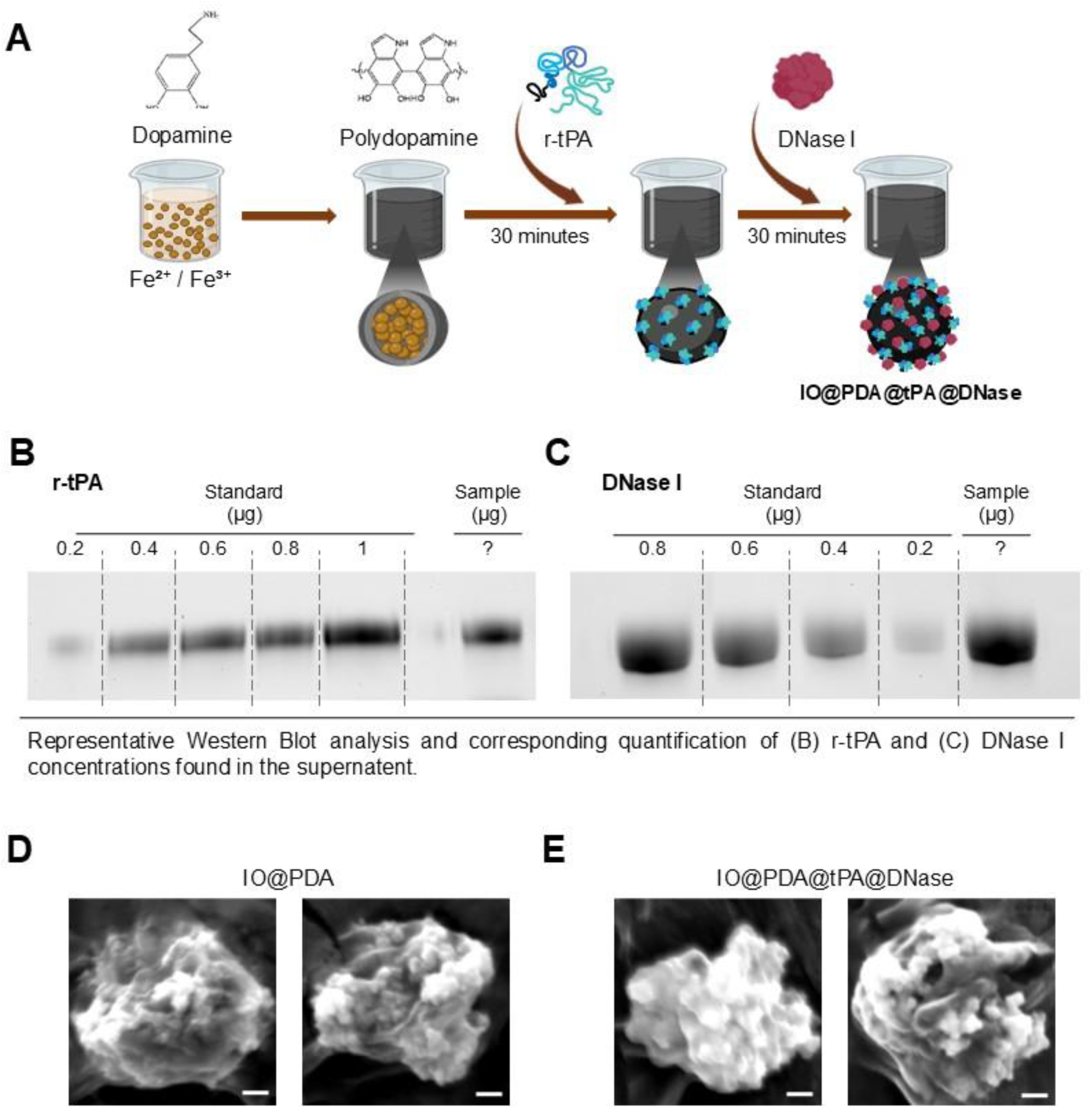
IO@PDA@tPA@DNase synthesis and characterization. **A.** Schematic illustration of the IO@PDA@tPA@DNase synthesis. Briefly, IO@PDA were first synthesized as described before^12^. Then, dialyzed r-tPA was mixed with the IO@PDA leading to the formation of the IO@PDA@tPA, then the DNase-1 was added leading to the new conjugate synthesis: the IO@PDA@tPA@DNase. **B-C.** Representative Western blot analysis and corresponding quantification of (B) r-tPA and (C) DNase I concentrations found in the supernatant. **D-E**. Scanning electron microscopy (SEM) micrographs of (D) bare IO@PDA and (E) the final IO@PDA@tPA@DNase conjugate. Scale bars = 100 µm.

**Table 1.** Summary of the IO@PDA@tPA@DNase properties.

|  | IO@PDA | IO@PDA@tPA@DNase |  |
| --- | --- | --- | --- |
| Hydrodynamic diameter (nm) ± SD | 614.6 ± 74.5 | 660.9 ± 56.3 | |
| Zeta potential (mV) ± SD | -18.63 ± 0.43 | -11.87 ± 0.427 | |
| r-tPA & DNase quantity fixed on the IO@PDA (mg/ml) | | **r-tPA**<br>0.55 ± 0.095 | **DNase**<br>0.317 ± 0096 |
| Iron doses injected to mice (mg[Fe<sup>2+</sup>]/kg) | 2 | 2 |  |
| r-tPA & DNase doses injected to mice (mg/kg) | 0 | **r-tPA**<br>2.5 | **DNase**<br>1.32 |

We then evaluated whether both enzymes retained their catalytic activity. DNA electrophoresis for DNase I activity verification revealed that the DNase I enzymatic properties remained intact even when grafted to the IO@PDA@tPA (Figure 2A). The Spectrofluor test, that allows to quantify tPA’s amydolytic capacities (Figure 2B, Figure S1), demonstrates that the r-tPA capacities are not influenced by the addition of DNase I to the conjugate (Figure 2B, Figure S1). The new conjugate, IO@PDA@tPA@DNase, was finally tested *in vitro* using human thrombi obtained post-thrombectomy from IS stroke patients (Figure 2C-F, Figure S2). Each clot was bisected to allow comparison between two treatment conditions using matched clot material (Figure 2C). Over two hours, clots were incubated with r-tPA vs IO@PDA@tPA (Figure 2C, D, Figure S2); r-tPA & DNase vs IO@PDA@tPA@DNase (Figure 2C, E, Figure S2); or IO@PDA@tPA vs IO@PDA@tPA@DNase (Figure 2C, F, Figure S2), allowing pairwise comparison. Quantification revealed no significant differences between the percentage of lysis for clots treated with r-tPA and IO@PDA@tPA (Figure 2D), indicating that conjugation of r-tPA to the particles does not impair its fibrinolytic activity compared to the free enzyme. The same was observed for clots treated with r-tPA & DNase vs IO@PDA@tPA@DNase (Figure 2E), suggesting that co-immobilization of both enzymes preserves their combined activity. However, the mean lysis percentage was higher, around 20% (Figure 2E), compared to the previous conditions (around 9%, Figure 2D), already highlighting the beneficial role of DNase on r-tPA-resistant thrombi. Finally, clots treated with IO@PDA@tPA@DNase were significantly more lysed, around 35%, compared to clots treated with IO@PDA@tPA (around 15%, Figure 2E), demonstrating the beneficial added effect of DNase to the conjugate through a synergistic effect. This synergy is even more evident when considering the final comparative analysis (Figure 2G).

**Figure 2.**
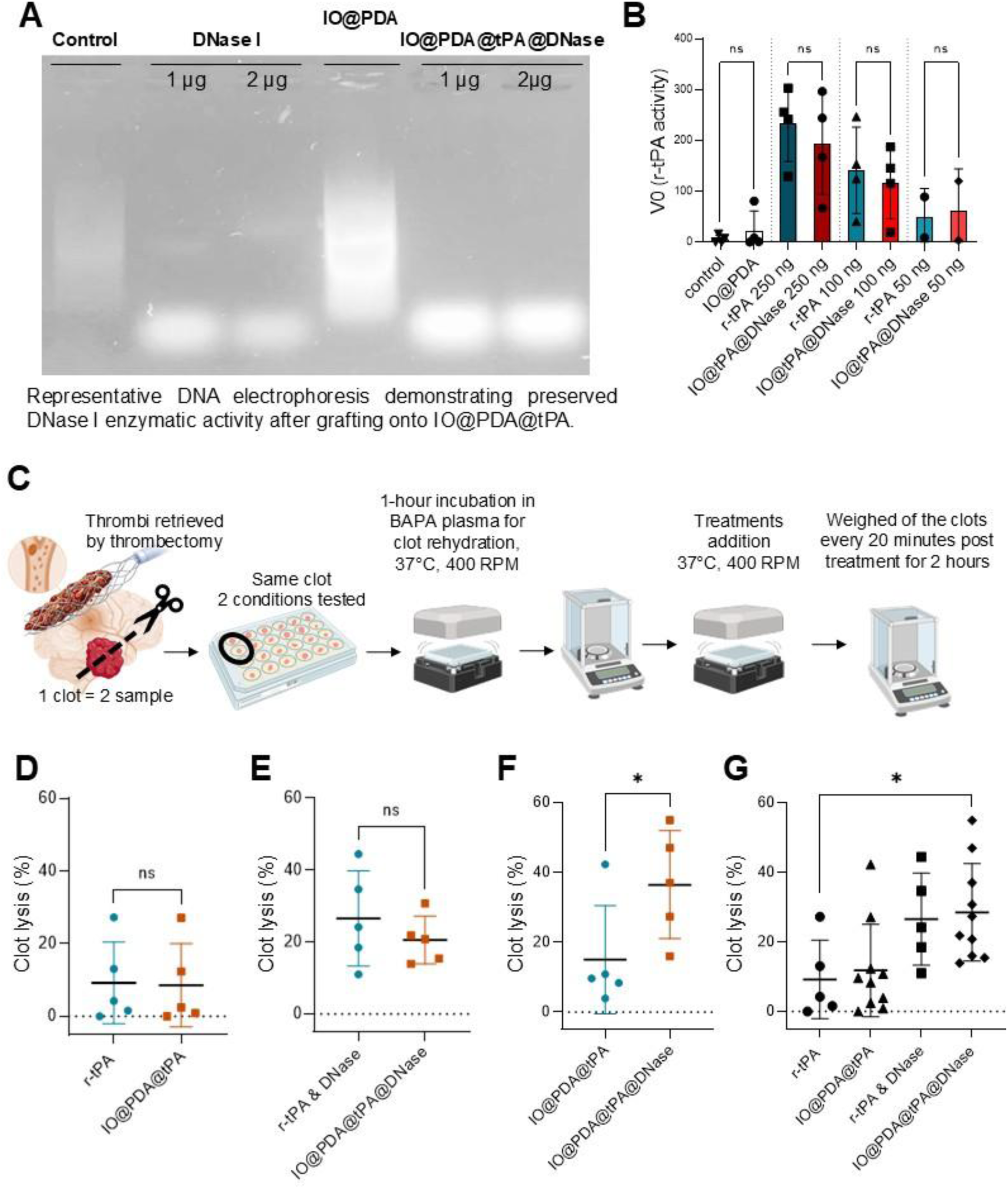
IO@PDA@tPA@DNase demonstrated superior lysis efficacy in vitro. **A.** Representative DNA electrophoresis demonstrating preserved DNase I enzymatic activity after grafting onto IO@PDA@tPA. **B.** Spectrofluorometric assay confirming that r-tPA amydolytic properties remain intact following the addition of DNase I to the conjugate. **C.** Schematic representation of the experimental design using human thrombi. **D–F.** Quantification of final clot lysis percentages for paired comparisons of clot weight loss over time: (D) free r-tPA vs. IO@PDA@tPA, (E) free r-tPA + DNase vs. IO@PDA@tPA@DNase, and (F) IO@PDA@tPA vs. IO@PDA@tPA@DNase. Data are presented as mean ± SEM. Statistical analysis was performed using one-way ANOVA with pairwise comparisons (n = 5). *p < 0.05, **p < 0.01. **G.** Mean clot lysis across experimental conditions, showing the superior efficacy of IO@PDA@tPA@DNase (Kruskal-Wallis’s test, multiple comparisons, *p < 0.05). Statistical analysis in this panel is provided for descriptive purposes only, as comparisons were not fully paired.

The therapeutic potential of the new conjugate was next assessed *in vivo*. The AlCl_3_-induced IS model was selected due to its ability to generate r-tPA–resistant clots^14^. Notably, this model does not induce microthrombi formation in the microvasculature^15^, precluding the use of molecular MRI for early-phase monitoring^13^. To overcome this limitation, functional ultrasound (fUS) imaging was employed to assess cerebral blood volume (CBV) before and during the acute phase of IS (Figure 3A). Five treatment groups were evaluated (Figure 3A-G, Figure S3): Saline; IO@PDA (2mg/kg); DNase (1.37 mg/kg); r-tPA (10 mg/kg); a combination of IO@PDA (2 mg/kg) + r-tPA (2.5 mg/kg) + DNase (1.37 mg/kg) and the IO@PDA@tPA@DNase conjugate (corresponding to 2.5 mg/kg r-tPA and 1.37 mg/kg DNase equivalency). Control groups (Saline, IO@PDA, DNase, r-tPA, and the free combination) all showed a sustained 60-80% CBV drop post-ischemia (Figure 3B-F; p<0.001, Figure S3). For mice treated with IO@PDA@tPA@DNase, a significant drop of CBV was indeed observed compared to baseline after IS as expected, however a rapid and progressive reperfusion occurred starting 10 minutes after treatment injection (Figure 3G, Figure S3) indicating effective revascularization.

**Figure 3.**
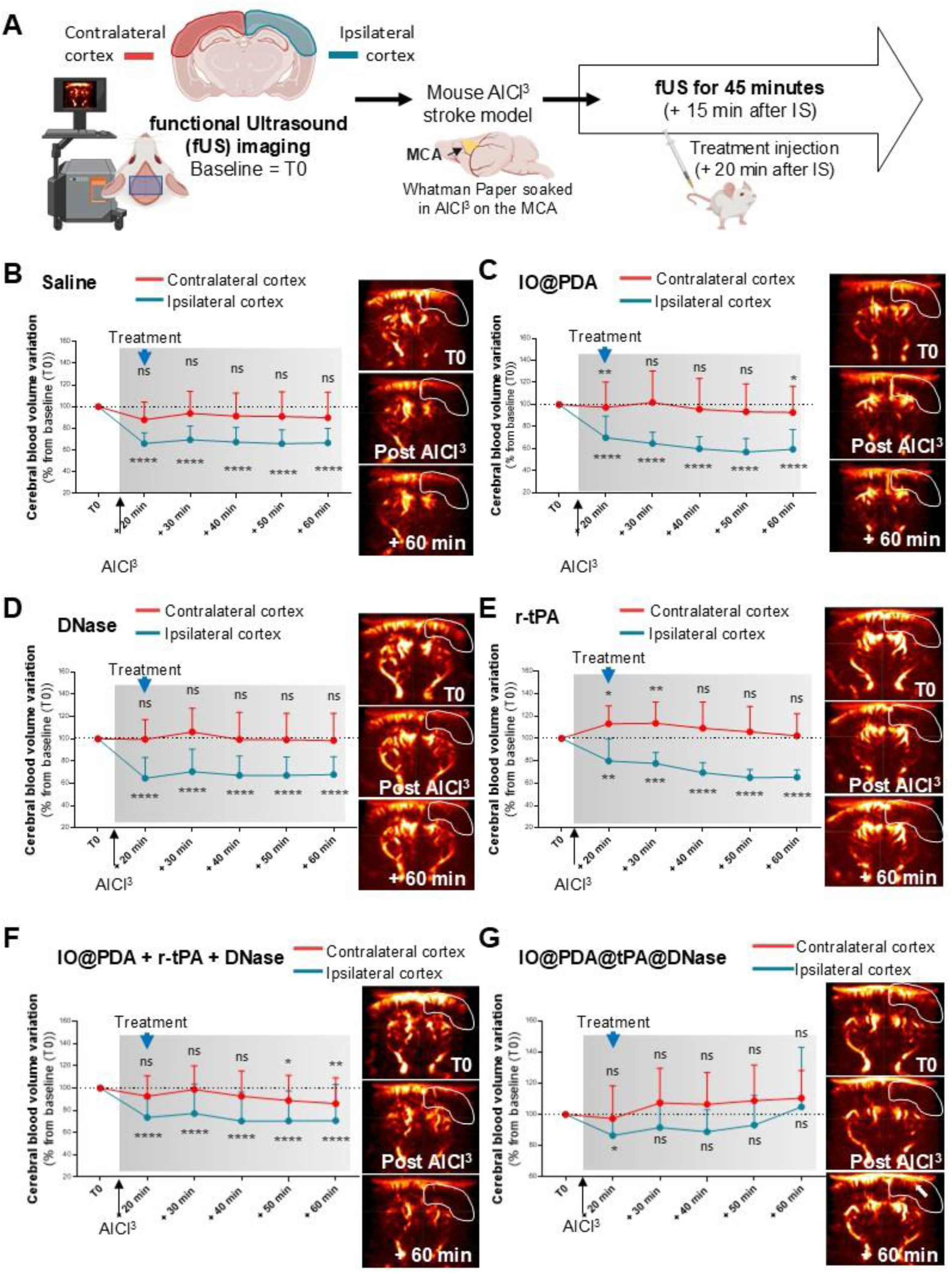
IO@PDA@tPA@DNase promotes early reperfusion. **A.** Schematic representation of the experimental design. Briefly, we used functional Ultrasound (fUS) imaging to quantify reperfusion in the ischemic area. Before inducing the IS, a 5-min acquisition was performed to have the baseline (T0) cerebral blood volume (CBV) of the mice. Then IS was induced by applying AlCl^3^ on the MCA for 5 minutes. 15 minutes after, a 45-minute acquisition began to follow any changes in the CBV in the ischemic area. 5 minutes after the beginning of the 45-minutes acquisition, one of the treatments: IO@PDA (2mg/kg); DNase (1.37 mg/kg); tPA (10 mg/kg); a combination of IO@PDA (2 mg/kg) + tPA (2.5 mg/kg) + DNase (1.37 mg/kg) and the IO@PDA@tPA@DNase conjugate (corresponding to 2.5 mg/kg tPA and 1.37 mg/kg DNase equivalency) was administered to the mice (corresponding to 20 minutes post ischemic events). **B.** CBV variation of the ipsilateral isocortex of IO@PDA-treated mice (percentage from baseline (T0), n=5, 2-way ANOVA, Multiple comparisons (T0 vs Tn+1), **** p < 0.0001). **C.** CBV variation of the ipsilateral isocortex DNase-treated mice (percentage from baseline (T0), n=6, 2-way ANOVA, Multiple comparisons (T0 vs Tn+1), **** p < 0.0001). **D.** CBV variation of the ipsilateral isocortex tPA-treated mice (percentage from baseline (T0), n=5, 2-way ANOVA, Multiple comparisons (T0 vs Tn+1), **** p < 0.0001). **E.** CBV variation of the ipsilateral isocortex IO@PDA + tPA + DNase-treated mice (percentage from baseline (T0), n=5, 2-way ANOVA, Multiple comparisons (T0 vs Tn+1), **** p < 0.0001). **F.** CBV variation of the ipsilateral isocortex IO@PDA@tPA@DNase-treated mice (percentage from baseline (T0), n=5, 2-way ANOVA, Multiple comparisons (T0 vs Tn+1), **** p < 0.0001). **G.** Representative images obtained during baseline (T0); 20 minutes and 60 minutes post IS for IO@PDA@tPA@DNase-treated mice.

To confirm the localization of IO@PDA@tPA@DNase at the thrombus interface, mice were euthanized 25 minutes after IS induction, corresponding to 5 minutes post-particle administration. For two-photon imaging, mice received an intravenous injection of lectin (green) prior to euthanasia to label the functional vasculature. Epifluorescence microscopy of brain sections stained for fibrinogen (magenta) and CD42b (grey) confirmed the presence of the thrombus (Figure 4A). Consistent with these findings, brightfield imaging revealed the accumulation of IO@PDA@tPA@DNase at the clot site (Figure 4A). This localization was further validated by two-photon microscopy and 3D reconstruction, which clearly showed the particles localized at the edge of the thrombus within the labelled vasculature (Figure 4B).

**Figure 4.**
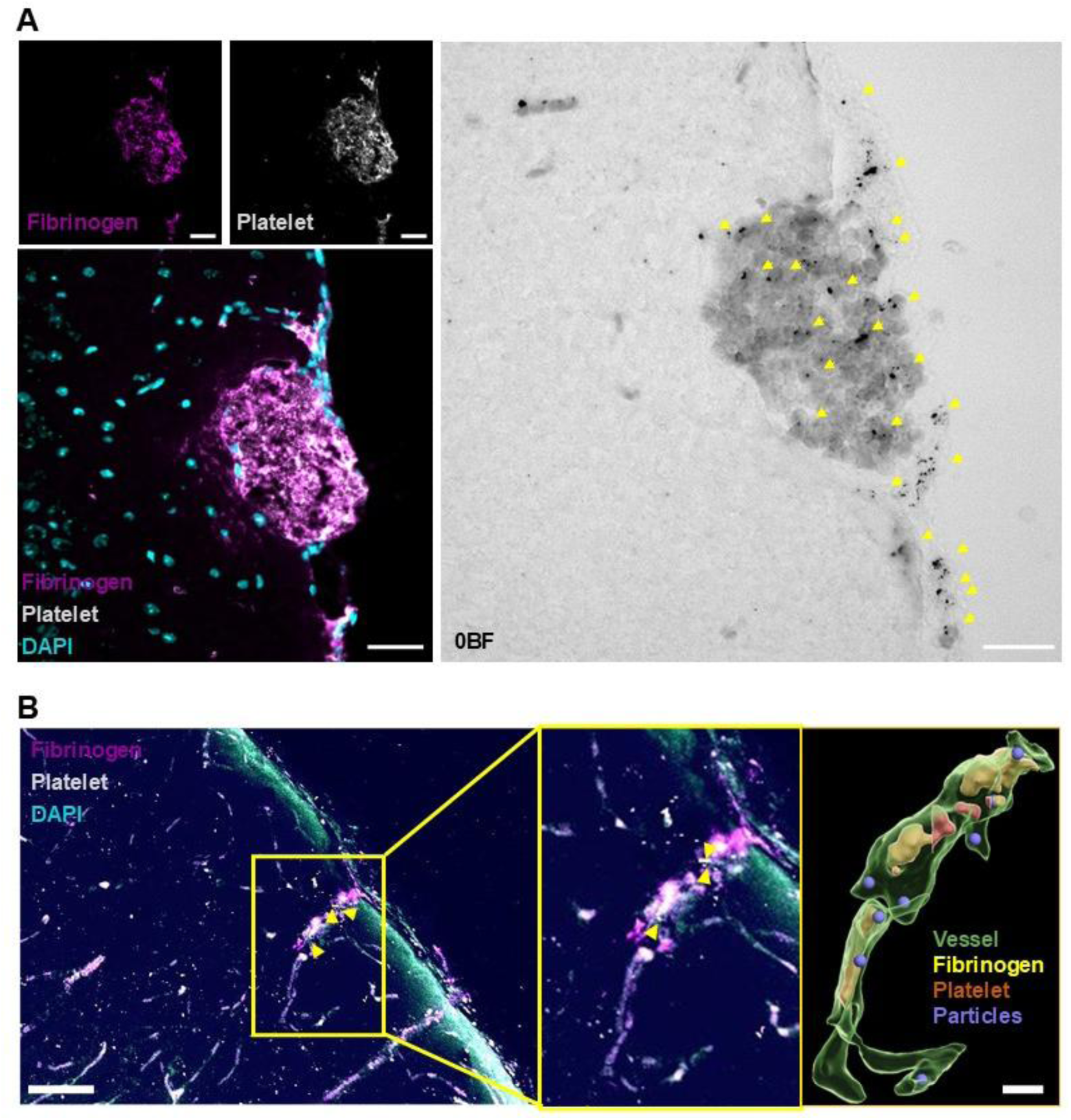
IO@PDA@tPA@DNase are found at the edge of the clot. **A.** Representative epifluorescence and brightfield micrographs of brain sections showing the thrombus composition: fibrinogen (magenta) and platelets/CD42b (grey). In the brightfield image, IO@PDA@tPA@DNase conjugates appear as black deposits localized within the clot (indicated by yellow arrows). Scale bar = 50 µm. **B.** Two-photon microscopy and 3D reconstruction illustrating the distribution of IO@PDA@tPA@DNase relative to the labeled functional vasculature (lectin, green) and the clot (fibrinogen and platelets). The 3D rendering highlights the accumulation of the nanoconjugates at the thrombus edge within the vessel lumen. Scale bar = 50 µm.

At 24 hours post IS, mice treated with IO@PDA@tPA@DNase exhibited a homogeneous reduction in lesion size compared to the other treatment groups which exhibited a heterogenous and larger lesion size (Figure 5A, B). Angiographic analysis performed 24 hours after IS revealed persistent occlusion in 63%, 67%, 58%, 86%, and 34% of mice treated with Saline, IO@PDA, DNase, r-tPA, and IO@PDA + r-tPA + DNase, respectively (Figure 5C, D). In contrast, 69% of mice treated with IO@PDA@tPA@DNase showed complete recanalization, further supporting the therapeutic efficacy of the conjugate (Figure 5C, D). Notably, the combined administration of IO@PDA + r-tPA + DNase also improved recanalization rates compared to single-agent treatments (Figure 5C). Finally, to assess functional recovery following IS, mice were subjected to the corridor test prior to IS induction, and at 1- and 5-days post-IS. Mice treated with Saline, IO@PDA, DNase, r-tPA, or the combination of IO@PDA + r-tPA + DNase showed impaired exploratory behavior at both day 1 and day 5 after stroke (Figure 5E, F; Figure S4). However, mice treated with IO@PDA@tPA@DNase presented no deficit (Figure 5F, Figure S4). In parallel, sensorimotor performance was assessed using the Corner test at baseline, and at 1- and 5-days following IS (Figure S4). At baseline, several groups (including Saline, IO@PDA, and r-tPA) already exhibited a moderate right-turning bias. Conversely, DNase, IO@PDA + r-tPA + DNase, and IO@PDA@tPA@DNase groups showed more balanced behavior (Figure S4). At 24 hours post-IS, all groups exhibited a marked rightward turning bias, reflecting acute sensorimotor deficits (Figure S4). Interestingly, only IO@PDA@tPA@DNase-treated mice maintained a symmetrical turning behavior, suggesting early functional recovery. By 5 days post-IS, most groups (Saline, IO@PDA, and IO@PDA@tPA@DNase) displayed near-symmetric turning patterns, indicating spontaneous or treatment-facilitated recovery (Figure S4). In contrast, mice treated with DNase alone continued to exhibit a strong right-turning bias, and those treated with r-tPA or the IO@PDA + r-tPA + DNase combination showed only partial improvement. These results further highlight the superior and sustained efficacy of the IO@PDA@tPA@DNase conjugate in promoting functional recovery.

**Figure 5.**
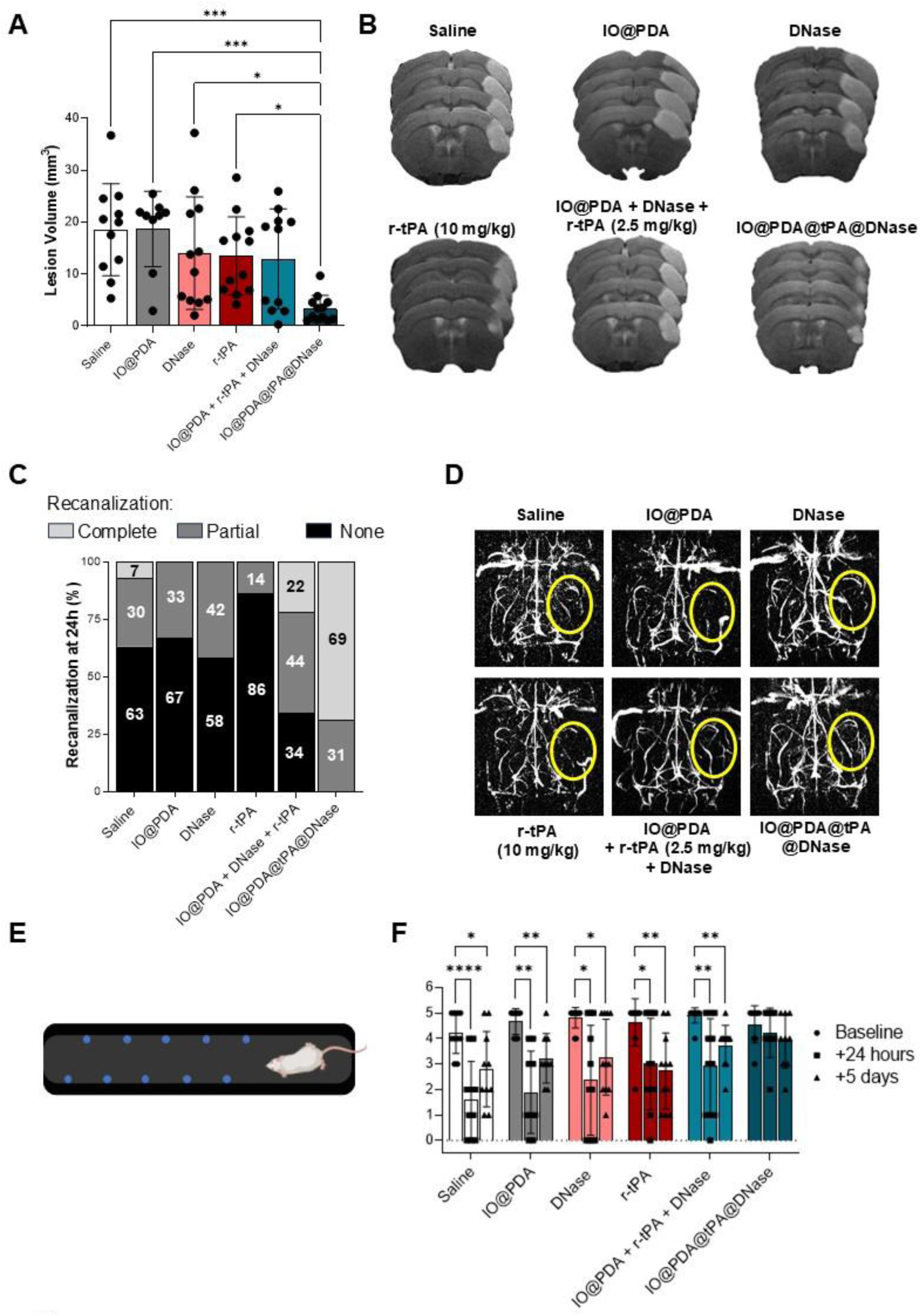
IO@PDA@tPA@DNase decreases the lesion size, promotes complete recanalization at 24 hours post ischemic event and improve functional recovery. A. **Lesion** volume (mm^3^) at 24 h after the injection of saline, IO@PDA, DNase, r-tPA (10 mg/kg), IO@PDA + r-tPA (2.5 mg/kg) + DNase or IO@PDA@tPA@DNase (One-way ANOVA with multiple comparisons (n = 9-12 per group). **B.** Illustrations of the lesion site area obtained after T*_2_*-weighted MRI. **C.** Angiography reveals a majority of non-recanalization of the MCA at 24h for mice treated with IO@PDA, DNase or r-tPA (10 mg/kg). IO@PDA + r-tPA + DNase appears to improve the recanalization at 24 hours but not as much as the IO@PDA@tPA@DNase-treated mice, for which a majority (75%) of complete recanalization was observed. **D.** Representative illustrations of the magnetic resonance angiography obtained for the mice depending on the treatment administered. **E.** Schematic representation of the corridor test apparatus (left). **F.** Quantification of left-side object visits by mice treated with Saline, IO@PDA, DNase, r-tPA (10 mg/kg), IO@PDA + r-tPA (2.5 mg/kg) + DNase, or IO@PDA@tPA@DNase. Assessments were performed before IS, and at 1 and 5 days post-IS (2-way ANOVA with multiple comparisons (n = 9-12 per group)).

## Discussion

The management of acute IS remains a major clinical challenge, primarily due to the narrow therapeutic window and the high rate of resistance to r-tPA. In this study, we developed and validated a dual-enzymatic nanoconjugate, IO@PDA@tPA@DNase, designed to overcome thrombus resistance by simultaneously targeting fibrin and NETs. This integrated approach significantly improves thrombolytic efficacy *in vitro* and promotes rapid reperfusion, reduced brain injury, and functional recovery *in vivo*.

Traditional thrombolysis focuses on fibrin, yet r-tPA-resistant clots are often rich in extracellular DNA and citrullinated histones—hallmarks of NETs that physically shield fibrin^6–8^. These NETs form a robust scaffold that traps platelets and red blood cells, physically shielding fibrin from r-tPA. Our *in vitro* results using human thrombi provide a strong proof-of-concept: while r-tPA or IO@PDA@tPA alone showed limited efficacy, grafting DNase I onto the nanoparticle dramatically increased lysis. This synergy suggests that DNase I de-compacts the clot matrix, increasing the fibrin surface area accessible to r-tPA. Notably, the conjugate outperformed the co-administration of free components, likely due to a “proximity effect.” By anchoring both enzymes on the polydopamine platform, we ensure simultaneous delivery and precise localization at the clot interface—confirmed by two-photon microscopy—preventing the dilution effect typical of systemic delivery.

However, the 35% lysis rate observed in human thrombi highlights remaining resistance, potentially due to von Willebrand factor or dense collagen cores. Furthermore, while the AlCl3-induced stroke model effectively produces r-tPA-resistant clots^14^, it triggers chemical *in situ* thrombosis rather than embolic occlusion. Consequently, the inflammatory profile and NET proportions may differ from clinical cardioembolic or atherosclerotic strokes. The absence of microthrombi in this model also precluded evaluation of the “no-reflow” phenomenon^15^; although fUS provided excellent macro-vascular temporal resolution, it lacks the cellular resolution to confirm capillary bed clearance.

Rapid recanalization in 69% of treated mice translated into significantly smaller, more homogeneous lesions and a complete absence of sensorimotor deficits in Corridor and Corner tests at 24 hours. This suggests that early, efficient revascularization is critical to preventing secondary neuroinflammation and permanent tissue loss.

A critical concern in any thrombolytic strategy is the risk of intracerebral hemorrhage. While our mice showed improved functional recovery and reduced lesion sizes, we must acknowledge that 20 minutes post-stroke is an early intervention point, which does not fully mimic the clinical reality where patients often arrive hours after symptom onset. As the BBB degrades over time, the risk of nanoparticles leaking into the parenchyma increases. Although our conjugate uses a relatively low dose of r-tPA, the potentiation by DNase I could theoretically increase the risk of hemorrhagic transformation if the BBB is severely compromised. Long-term safety studies focusing on the late therapeutic window (3 to 6 hours post-stroke) and the systemic clearance of iron oxide particles are essential before clinical translation can be considered. From a translational perspective, the hydrodynamic diameter of our conjugate (∼660 nm) is relatively large for nanomedicine standards. While this size favors accumulation at the edge of a large occlusion due to flow dynamics, it might limit the penetration of the particles into smaller collateral vessels or highly constricted areas. Additionally, the 72-hour incubation required for primary antibodies in our histological analysis underscores the density of these tissues; it remains to be seen how effectively these large conjugates can diffuse into the core of an established, retracted human thrombus in a high-pressure arterial environment.

## Acknowledgements

AI was used only for grammatical and orthographic correction in the text. Some elements in the figures were generated via BioRender.com.

## Authorship Contributions

A.P. designed the study, performed the experiments, analyzed the data, and wrote the manuscript. M.L., C.H., A.T., I.K., performed experiments and data analyses. S.d.M.L. and M.G. designed the nanoparticles (IO@PDA). M.S.N., B.H.T.N provided access to the thrombectomised clot biobank and helped with clot lysis experiments. D.G. performed SEM acquisition. D.V. and T.B. contributed to the project conception and manuscript writing.

## Supplemental Data

**Figure S1.**
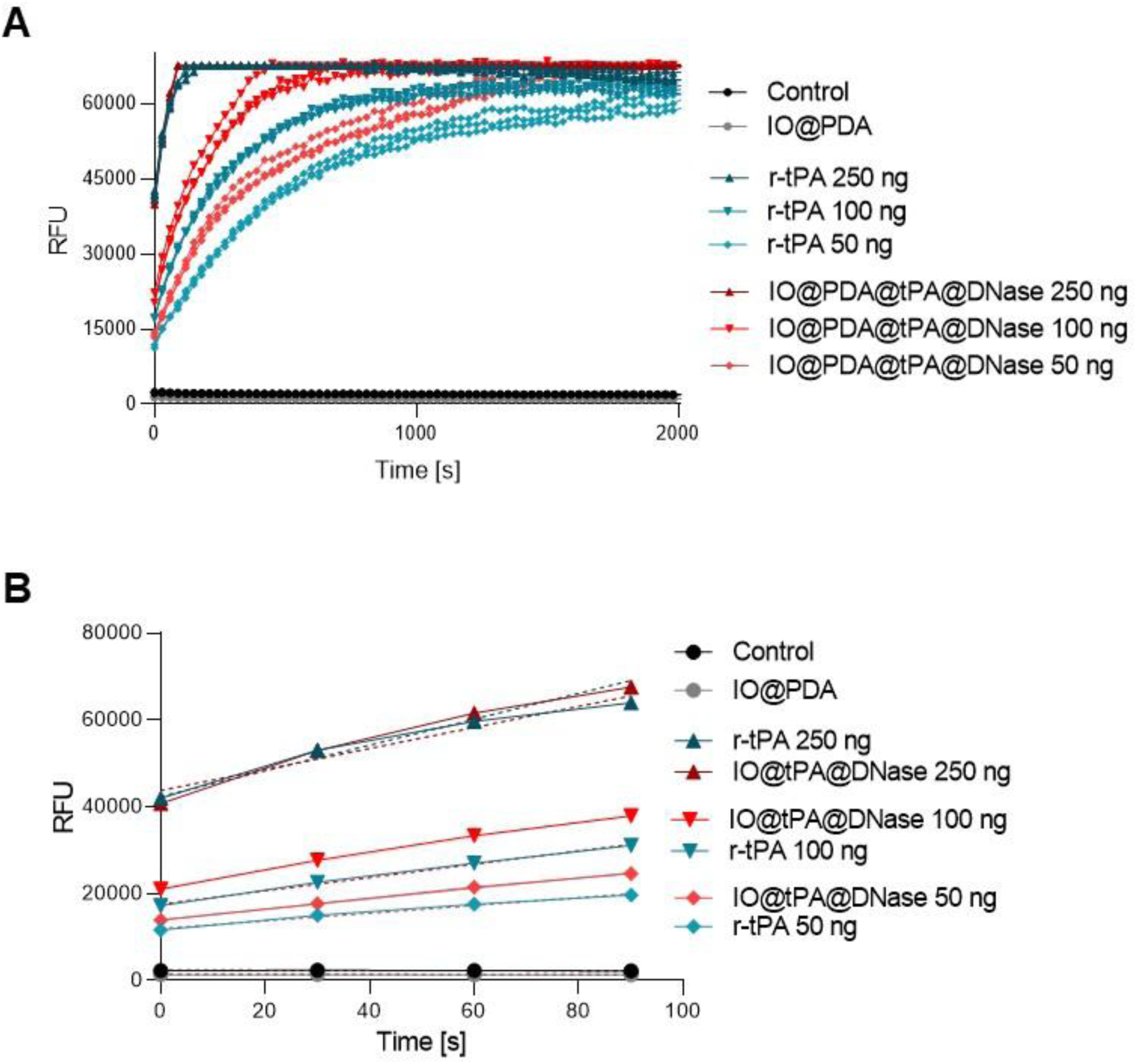
***S*pectrofluorometric assay. A-B.** Representative curves (A) and slopes (B) obtained after the spectrofluorometric assay.

**Figure S2.**
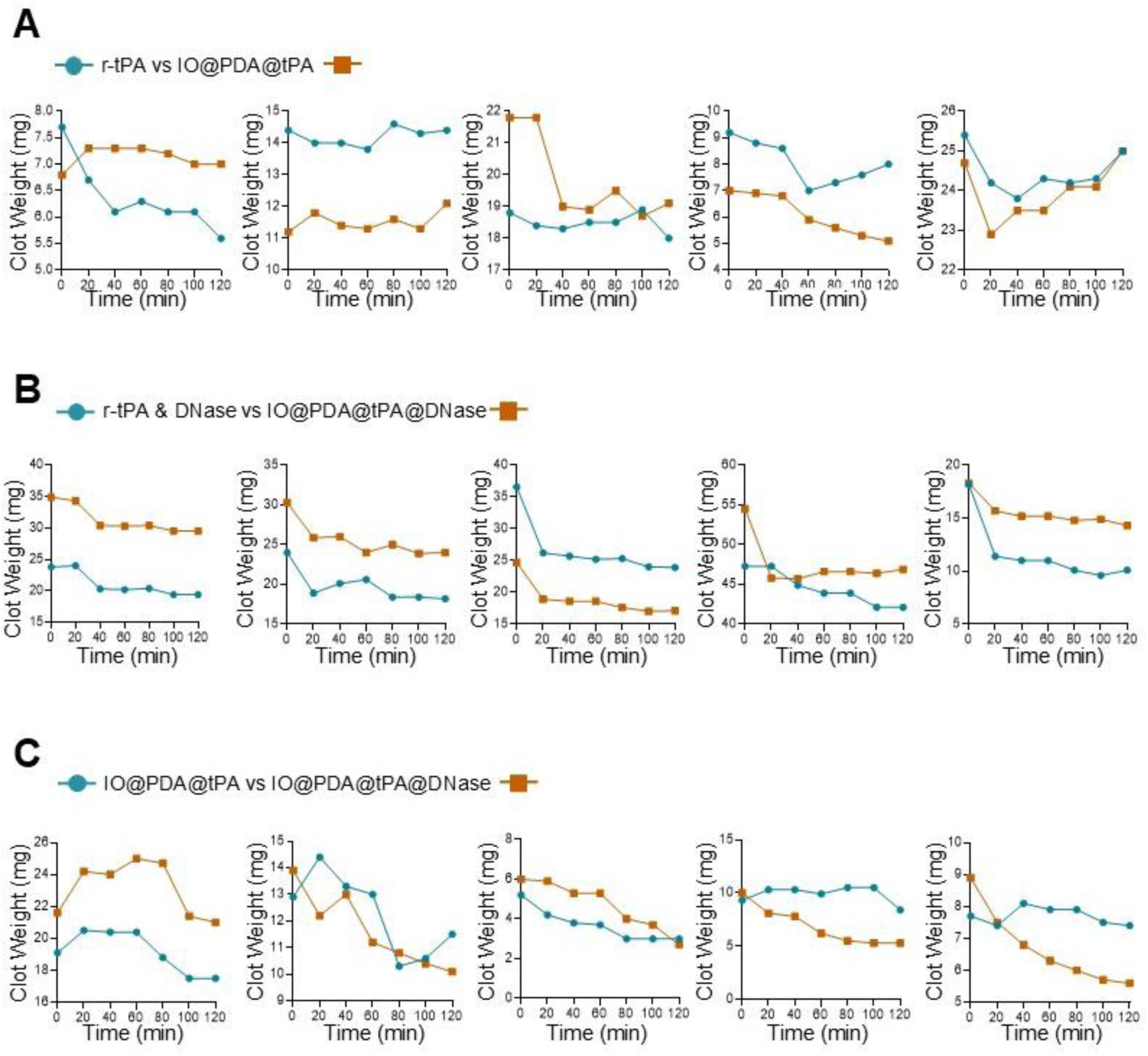
Clot weight loss over time for individual paired comparisons. **A**. Free tPA vs IO@PDA@tPA. **B.** Free tPA + DNase vs IO@PDA@tPA@DNase. **C.** IO@PDA@tPA vs IO@PDA@tPA@DNase.

**Figure S3.**
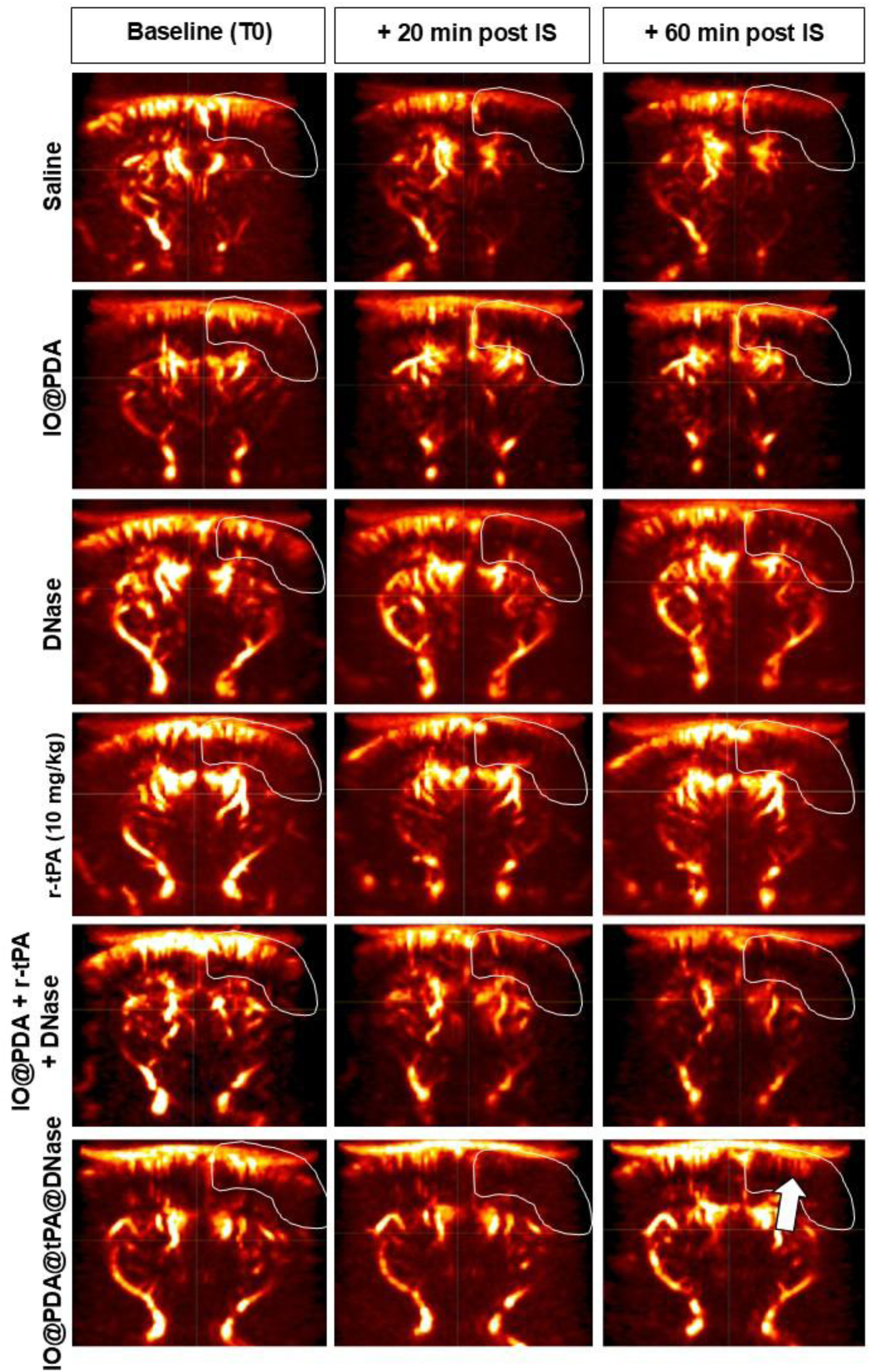
Illustration of fUS imaging of cerebral perfusion. Representative fUS maps obtain showing power doppler intensity for all the experimental groups tested at Baseline (pre-ischemia), 20 minutes and 60 minutes post IS induction.

**Figure S4.**
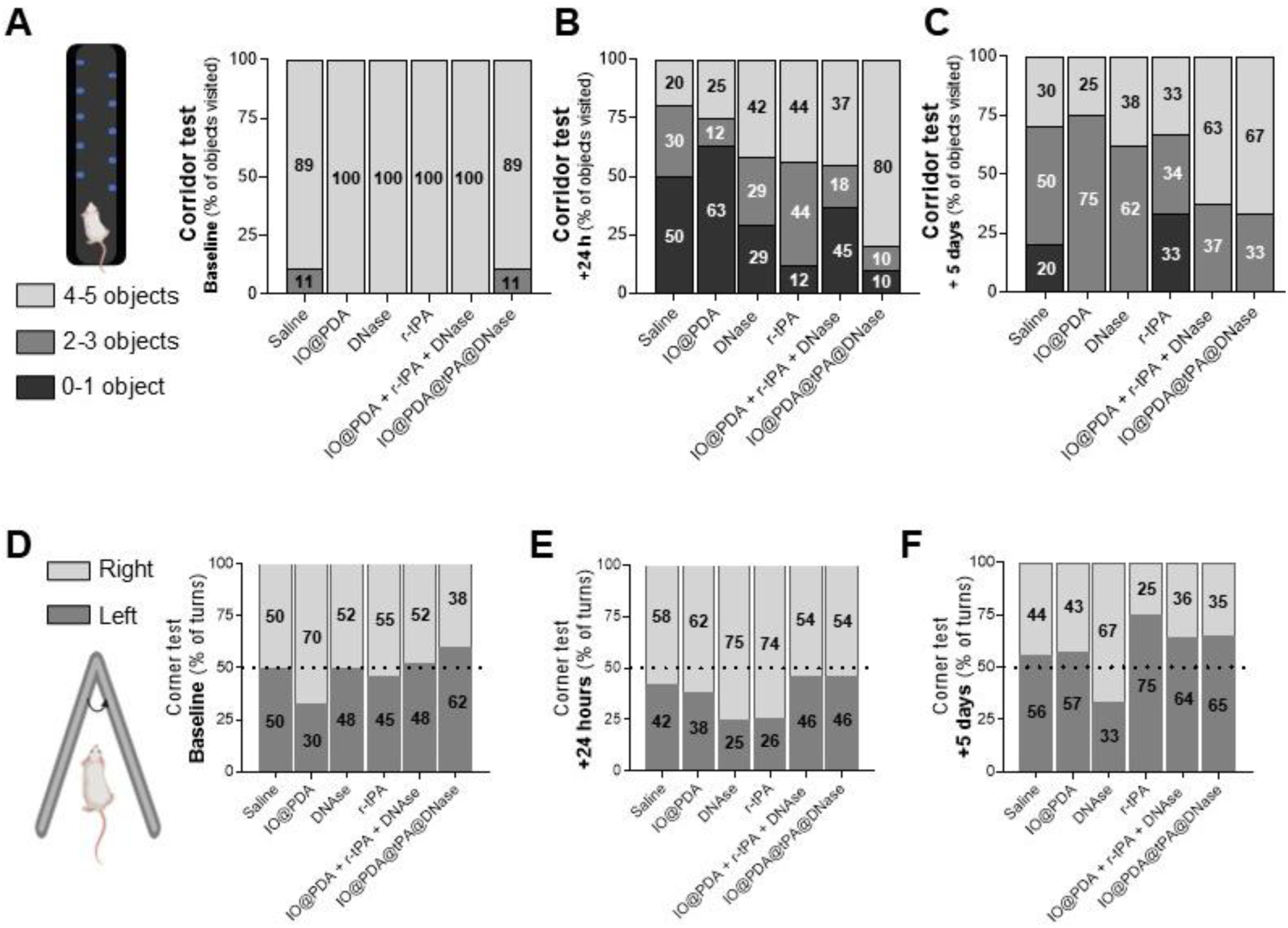
Functional recovery. A-C. Detailed analysis of the proportion of left-side object visits for each group at baseline (A), 1 day (B), and 5 days (C) after IS (n = 9-12 per group). **D-F.** Evaluation of motor and sensorimotor function using the corner test. Percentage of turns to the left or right before IS (D), at 1 day (E), and 5 days (F) post-IS (n = 9-12 per group).

